# Impact of the Microtubule Cytoskeleton on Insulin Transport in Beta Cells: A 3D Computational Study

**DOI:** 10.1101/2025.02.12.637971

**Authors:** Thoa Thieu, William R. Holmes

## Abstract

Glucose-stimulated insulin secretion (GSIS) in pancreatic *β* cells is vital to metabolic homeostasis. Recent evidence has highlighted the critical role of the cells’ microtubule (MT) cytoskeleton in regulating transport and availability of insulin containing vesicles. How these vesicles move within the cell and how that mobility is influenced by the MT network is however not well understood. The MT network in these cells is dense and randomly oriented. Further insulin vesicles are relatively large compared to the spaces in this dense meshwork. Here we develop a 3D computational model that simulates vesicle motions in the dense MT network of the *β* cell. The structure of this MT network, along with the dynamics of vesicle motions, are calibrated to microscopy data from *β* cells to ensure physiological relevance. Our results reveal a number of key observations. 1) The MT network in *β* cells likely impairs motion of larger vesicles (200 − 300*nm* in diameter). 2) This is in part a consequence of their “caging” by the MT network. 3) This results in a substantial reduction in the likelihood of vesicles transiting from the cells interior to the plasma membrane, a pre-cursor to GSIS. 4) Dynamic remodeling of the MT network reduces the strength of these effects. 5) That same remodeling however introduces anomalous (sub-diffusion) motion characteristics. Taken together, these results indicate that the dense MT network of the *β* cell substantially inhibits mobility and availability (for GSIS) of insulin. It further sheds light on how the complex filament network in cells leads to statistically anomalous motions. Finally, this modeling further provides a test-bed for determining how potential manipulations of the structure and dynamics of this network would tune GSIS.

**SIGNIFICANCE:** Insulin release from pancreatic *β* cells is crucial for blood sugar regulation, and recent research suggests the microtubule network inside these cells plays a key role in how insulin is transported and released. This study developed a 3D computational model to explore how insulin vesicles move through this dense network. Results show that the microtubules can “cage” larger vesicles, making it difficult for them to reach the cell surface for insulin release. Dynamic remodeling of the network can however increase insulin mobility and availability. These findings highlight the impact of the microtubule network on insulin transport and secretion and provide insight into potential ways to tune this process.

## INTRODUCTION

Insulin is a critical hormone and its dysregulation can affect energy balance and lead to metabolic disorders including diabetes, which itself effects ~ 9% of the population in the USA (1–7). Understanding its regulation is thus critical to modeling metabolic homeostasis and the onset such disorders. Regulation of insulin can be investigated at the systemic level (e.g. systemic hormone balance), the organ level (the pancreas or the Islets that comprise it), or the cellular level. Here we investigate the mechanisms that regulate insulin availability and release at the level of individual pancreatic *β* cells.

*β* cells are the primary source of insulin production in the body. In these cells, the endoplasmic reticulum (ER) and the Golgi package insulin protein into vesicles (also referred to as “granules”) that are distributed throughout the cell (8). Upon stimulation by glucose, cells exostosis these vesicles in a process referred to as Glucose-Stimulated Insulin Secretion (GSIS) (9). The regulation of GSIS is crucial for maintaining balanced blood glucose levels, and tightly controlled release of insulin is essential to prevent hypo- or hyperglycemia. A key determinant of how much insulin is released during GSIS is the size of the pool of “readily releasable” vesicles—those that are both close enough to the plasma membrane to be released and biochemically capable of docking to it (10). Here we will investigate the mobility and localization of insulin, but not biochemical modifications that facilitate docking.

The structure and function of the cells’ cytoskeleton has been found to play a vital role in secretion (11–13). Microtubule (MT) filaments and the motors that walk on them are a major cellular transport system (14, 15) in many contexts. This transport is known to effect *β* cell function (16–18). Glucose stimulation enhances MT-based directed motion of vesicles, a crucial step in insulin secretion (19). Advanced glycation end products (AGEs) impair GSIS by disturbing the MT cytoskeleton in *β*-cells, with p38/MAPK activation playing a pivotal role in MT depolymerization (20). Recent work (13) has demonstrated that short term depletion of MTs resulted in increased secretion, consistent with earlier findings (21, 22).

Our prior modeling (23) indicated that one of the main functions of the MT network is to inhibit the avaliability of readily releasable insulin. Modeling combined with subsequent super-resolution image quantification demonstrated there is a sub-membrane network of MT filaments that bind and remove insulin from near-membrane region, rendering it unavailable. Depolymerization (nocodazole for example) or destabilization (by glucose stimulation) of these filaments both lead to increased secretion (13). Further, selective stabilization of this sub-membrane network by Tau protein (embedded in the membrane) slows secretion by specifically reducing the density of insulin near the membrane (24).

Despite the extensive studies on how MT dynamics influence insulin secretion, a key aspect of vesicle transport in *β*-cells remains under-explored. How do they move within the complex environment of the cell and does the MT network influence that mobility? Insulin vesicles are relatively large compared to the density of the network they reside in and the network itself is randomly oriented in the cells’ interior (13, 23). Estimates suggest there are on the order of 10,000 (25) vesicles in *β* cells with sizes 100-300nm in diameter (26). These vesicles are thus roughly the same size as the spaces between filaments in the MT network: 70% of such spaces are < 240nm in diameter (13). Additionally, single particle tracking has demonstrated that insulin vesicle motions are anomalous (13, 27) and are neither diffusive nor ballistic. This behavior is frequently observed in biological systems with complex, heterogeneous environments, such as the intracellular cytoskeleton (27–31). In the case of insulin, periods of intermittent mobility are clearly visible in the vesicle tracks (13) and the statistical patterns of immobility obey a particular type of anomalous motion referred to as a Continuous Time Random Walk (CTRW) (27).

Taken together, this evidence leads to a hypothesis that the density of MT filaments relative to the size of vesicles impairs their motion. This can occur due to two possible mechanisms: 1) the physical presence of MTs reduces the effective speed of motions and 2) the filament network can physically cage vesicles and sequester them in place. In this study, we construct a computational model of the motions of insulin vesicles constrained by the MT filament network to assess the validity and consequences of these effects. To ensure the validity of these results in physiological conditions, properties of the MT filament network and the vesicle motions within it are calibrated to microscopy data where possible.

In this work, we investigate the complex dynamics of insulin vesicle transport within pancreatic *β*-cells, and how the MT network constrains those dynamics. We will focus on both practical and theoretical consequences of this vesicle-MT network interaction. On the practical side, we will assess how vesicle-MT interactions influence the likelihood of vesicles reaching the membrane. Reaching the membrane is a required pre-cursor to secretion and thus critical to proper function. On the theoretical side, we will assess how vesicle-MT interactions influence the statistical properties (time averaged mean squared displacement specifically) of motions. Here we are specifically interested in understanding to what extent these interactions may be the source of the anomalous motions observed in (13, 27). We will specifically focus on the influence of three aspects of this system that influence vesicle-MT interactions: 1) vesicle size, 2) MT filament density, and 3) MT dynamics.

Results demonstrate that the interplay between vesicle size and MT filament density significantly influences vesicle mobility. Further, the nominal density of MTs observed in *β* cells likely significantly restricts vesicle mobility and substantially reduces the likelihood of vesicles interacting with the cell membrane. The vesicle - MT interactions are however not sufficient to explain previously observed anomalous motions. Instead, the “caging” observed in vesicle tracks may be a consequence of the dynamic nature of the filament network. Taken together, these results indicate that both the density and dynamics of the MT network influence vesicle mobility and availability for functional secretion.

## METHODS

### Overview

In this discussion, we outline the construction of a 3D computational model for insulin-producing *β* cells in the pancreas, inspired by the previous findings of (13, 27) and the prior modeling in (23). Our methodology has two essential components. First, we construct a MT filament network that mimics the statistical structure of MT networks in *β* cells. Second, we simulate random vesical motions in the cell constrained by the steric (physical) interactions between the vesicle and the filaments. These simulations are used to determine how the filament network restricts the mobility and availability of these vesicles and how this is influenced by 1) vesicle size, 2) MT network density and 3) the dynamics of the network.

### MT Construction

We approximately model the cell as a sphere in 3D with a diameter of 10*μm*. Unlike other cells where MTs are nucleated from a MT organizing center (MTOC), MTs in *β* cells are nucleated from the golgi (32) and form a randomly oriented network of filaments (13, 23). We thus construct a network of randomly located and oriented filaments in the *in silico* cell. For computational tractability, we make a few simplifying assumptions. 1) Each filament is straight. 2) Filaments do not interact with each other. They interact with the simulated vesicle, but not each other. 3) Filaments are fixed in their location and length. They do not move within the cell. Further, they do not undergo dynamic instability, which is less prevalent in *β* cells than other cell types (13). 4) Each filament that interacts with the cell membrane / perimeter simply terminates at the membrane. We do not account for bending of filaments by the membrane. Here we will be primarily interested in the motions of vesicles in the cells’ interior and thus do not construct a sub-membrane array (23). 5) We assume every filament can de-polymerize completely (with new filaments created to maintain density). This will be modeled as the complete disappearance of the filament (and commensurate appearance of a new one). We do not explicitly model the (de)polymerization of each filament over time. The resulting network of MTs is a random collection of filaments, each fixed in their location, but where filaments can randomly appear and disappear.

The essential question we will investigate is how vesicles move in this network. Toward this end, we construct a network that mimics the basic statistical structure of MTs in *β* cells. We do not have access to number and length distributions of filaments in *β* cells. However it was found (13) that ~ 65-70% of the spaces within the MT network were <240nm in size. This is important since the typical size of an insulin vesicle is 200-300nm in diameter. This serves as a proxy for MT density that we will use to calibrate the MT network. To mimic this density, we determine a combination of number (*N*) and length (*L*) of filaments that yield a network with this property.

For each combination of (N,L), we construct a network. Within this network, we seed a 1,000 random points and determine the size of the space near that point. See the “Space Size Determination” section for how the sizes of these spaces are computed. With these 1,000 “space” sizes we compute the fraction that are <240nm in diameter. Results of this analysis are shown in Table 1. The combination (N=15,000, L = 7) generates a network with 68% of spaces <240nm. We will use this combination as representative of a *β* cell. The combination (N=10,000, L=4) produces a network with larger spaces. We will refer to these as “dense” and “sparse” networks throughout this article. Multiple networks with different combinations of (N,L) will exhibit this ~65-70% of spaces <240nm. For simplicity, we use only these two combinations for further analysis.

**Table 1:** Table illustrating the fraction of spaces in the filament network that are <240nm as a function of the number (N) and length (L) of filaments. For each N, L, we constructed a random network of filaments. We then seeded 1,000 points, and for each point found the largest sphere that could be inscribed near that point without intersecting a MT. We then quantify the fraction of these spheres that have a diameter <240nm. The ^**^ value (L=7, N=15,000) is the set of values that we will use to construct a network statistically mimicking the data in (13). The ^*^ value will be used as an example of a network with a larger space distribution. These will be referred to as “dense, sparse” networks respectively throughout.

| $L \backslash N$ | 10,000 | 12,500 | 15,000 | 17,500 | 20,000 |
| --- | --- | --- | --- | --- | --- |
| 4 $\mu\text{m}$ | 0.34* | 0.47 | 0.54 | 0.62 | 0.71 |
| 5 $\mu\text{m}$ | 0.41 | 0.53 | 0.64 | 0.69 | 0.78 |
| 6 $\mu\text{m}$ | 0.43 | 0.56 | 0.65 | 0.75 | 0.82 |
| 7 $\mu\text{m}$ | 0.44 | 0.57 | 0.68** | 0.77 | 0.82 |
| 8 $\mu\text{m}$ | 0.44 | 0.60 | 0.70 | 0.79 | 0.84 |

#### Filament dynamics

MT filaments in *β* cells are mostly stable in the sense that they do not undergo frequent dynamic instability. They do however exhibit dynamic properties. Filaments can destructively de-polymerize and new filaments can be nucleated. In non-perturbed cells, these processes are in balance so that the statistics of the network structure are consistent over time (13). To account for this dynamic, we assume each MT has a half-life *τ*_*half*_ = 10, 100, 1, 000, 10, 000, ∞ sec. These represent a wide range of filament dynamic timescales from fast to completely static.

To describe these dynamics, at each simulation time step, we will model the number of catastrophe events in that time step as Poisson distributed with the relevant half-life. Each filament that is destroyed will be replaced with a new filament at a random location that does not overlap the position of a vesicle. Other forms of dynamics such as filament bending and transport (via motors) are present. However incorporating these would drastically increase computational complexity. Here we use this catastrophe / nucleate dynamic as a proxy for general dynamics to determine the effect of such dynamics on vesicle mobility.

#### Space size determination

In the previous section, we describe how to construct a filament network whose space size distribution mimics that of *β* cells. Here, we describe how the sizes of spaces are computationally determined for *in silico* networks. We define a space in the network as the largest sphere that can be circumscribed in a local region of the cell without intersecting any filament. The diameter of this sphere is then the “size” of the space.

The following is our basic approach to determining the size of a local space. Randomly choose any point in the cell (at least 1 *μ*m from the membrane to avoid membrane interference). Conceptually, think about expanding a spherical balloon from that point, allowing the balloon to move in space until it can no longer expand. The diameter of the resulting balloon is the size of the space locally near that seed point. We will implement this conceptual approach computationally.

First, define the shortest distance between a point and a line segment as follows. Let the point be **p** ∈ ℝ^3^. The line segment is defined by the start point **s** ∈ ℝ^3^ and the end point **e** ∈ ℝ^3^. The general formula for the distance *d* from the point at **p** to the MTs defined by segment **s** to **e** is:

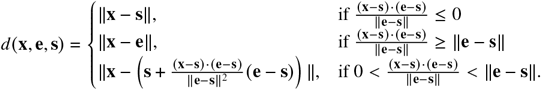

This formula encapsulates all scenarios for calculating the shortest distance from the granule to the MTs.

We use this calculation as the foundation of an optimization process to iteratively move a seed point in space until it is at the center of the largest possible sphere that can be circumscribed in that local area of the network. Begin with a seed point **x**_0_ = (*x*_0_, *y*_0_, *z*_0_). Define a cost function as

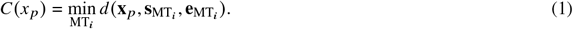

For the maximum size circumscribed sphere in a region near **x**_0_, *C* will be maximized. Further, the *radius* of the resulting sphere will be the maximal value of *C*. The size of the space near **x**_0_ can then be determined by computing the maximal value of *C* as

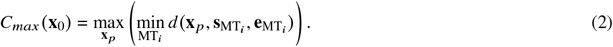

Figure 1 illustrates this approach visually. The figure shows the value of *C* as a function of position in a simple configuration of MTs.

**Figure 1.**
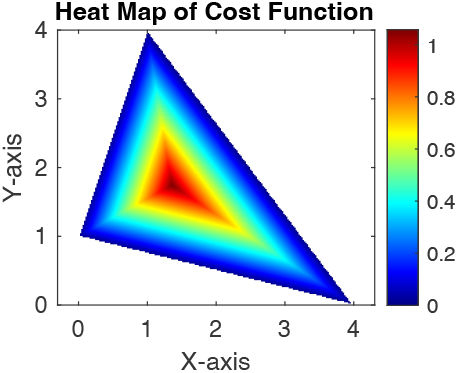
(Color online) 2D schematic of the cost function used (Equ. (1)) used to determine space sizes. The three sides of the triangle depict MT filaments in 2D and the heatmap indicates the value of the cost function within the triangle. The peak of the cost function represents the center of the largest inscribible circle and the peak value represents the radius of that circle.

To compute this optimum for **x**_0_, we use gradient descent on − *C* (**x**_**p**_) with **x**_0_ as the initial state of the optimization. It is critical that a true “local” optimizer is used for this. A non-local optmizer may allow the optimization to jump over a filament into a new space. For each seed point, we want to find the size of the space near that seed point. This effect of jumping to a new space would skew space size distributions. Simple gradient descent with no additions (not momentum, Ada-grad, ADAM, or any extension) is used with an appropriately small iteration step size to ensure this does not occur.

### Network constrained transport

In real *β* cells, vesicle motions are motor (both MT and actin) driven. However, observed motions appear diffusive (according to mean squared displacement) (13, 27), likely due to the presence of multiple motors on each vesicle interacting with different filaments. As a result, we do not consider individual motor motions and instead model vesicle motions as diffusive but constrained by the network of filaments. We note that vesicle motions have been previously been modeled as sub-diffusive (23, 27, 33). However this characteristic is hypothesized to be a consequence of the interaction between vesicles and their environment, which is the topic of this work. Thus we model the free-space motions as standard diffusion and assess how interactions with MTs augment those motions.

For simulations, we use a lattice-based random walk modified to accomodate the filaments as obstacles that prevent steps on the lattice. Following (23), we set the diffusion rate to be *D* = 0.01*μm*^2^/*s*. The lattice step size is fixed at 20nm and the time step is chosen according to 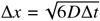, resulting in Δ*t* = 6.7ms. At each time step, a random integer (1,2,3) is chosen to determine which dimension (x,y,z) the step will be taken and a second random integer (−1,+1) is chosen to determine which direction the step is in that dimension. This is a standard lattice random walk algorithm that, in the absence of obstructions, yields a random walk with mean squared displacement that scales appropriately with the chosen diffusion constant.

Vesicles of different size will be considered. Note however that we do not modulate the diffusion coefficient by the vesicle size. While vesicle motions are statistically diffusive (rather than ballistic), those motions are motor driven and so the standard Einstein relation for a passive diffusion coefficient is not appropriate. Further, we do not have data in this system for diffusion rate as a function of vesicle size. For simplicity we thus fix the diffusion coefficient.

This random walk is modified by the presence of filaments as obstructions. To account for this, the radius of the vesicle (*r*) is needed. When a random walk step is attempted, if the distance between the vesicle location and *any* filament is < *r*, that step is aborted and the time step increments forward.

The motion of each vesicle is simulated for 60min of time and the trajectory is recorded. For every unique simulation condition, we will simulate the motion of 200 identical vesicles with different initial locations for statistical purposes. These are 200 separate and independent simulations; the vesicles do not interact with each other. For simulation analysis, we quantify whether the vesicle is able to reach the membrane (defined as coming within 200nm of the membrane) and the mobility properties. These will be quantified as a function of 1) the size of the vesicle, 2) the density of the network, and 3) the speed of filament network remodeling (filament half-life).

#### Vesicle path analysis

The central goal of this work is to determine how the filament network influences mobility and availability of vesicles. We define “availability” as the potential for a vesicle to reach the membrane within the 60min simulation time. Note that the primary and secondary secretion responses in glucose stimulated *β* cells are ~ 5, 60 minutes respectively (34). Thus, for each simulation condition (200 identical simulations), we will quantify the fraction that reach anywhere within 200nm of the cell membrane (since membrane docking proteins can extend a short distance to dock vesicles).

Quantifying properties of granule mobility is more complex since there are multiple facets of mobility. Following (27), we will quantify Time Averaged Mean Squared Displacement (TAMSD) of each trajectory

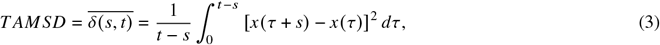

where *x* is the position of the vesicle, *s* denotes the lag time, and *t* represents the total measurement time for each trajectory. For simple diffusion, this quantity should be log-linear with respect to *s* with slope 1. Any deviation from this represents a deviation from simple diffusion. Additionally, TAMSD should be constant with respect to *t* for diffusion. A decrease in TAMSD as a function of *t* is indicative of an “aeging” phenomenon, which Tabei et al. (27) observed in insulin vesicle tracks. TAMSD will be computed and plotted with respect to both *s, t* for analysis of deviations from simple diffusion.

We will also quantify the tendency for vesicles under different conditions to be spatially constrained or trapped by the network. For an individual trajectory, this will be computed as follows. We quantify the maximum duration during which the vesicle does not move more than 300nm in displacement. Each trajectory is an ordered list of locations (*x*_1_, *x*2, …, *x*_*n*_) where *t*_*i*_ = *i* * Δ*t* is the time of each observation. For each *i*, define *s* = *x*_*i*_ as the starting location from which we look forward in time. Find the first *j* such that *x*_*j*_ − *s* > 0.3 and define *τ*_*i*_ = *t* _*j*_ − *t*_*i*_. This is the time it takes the particle to move 300nm from the anchor location *s*. Now define *τ* = max {*τ*_*i*_} as the maximum of such times. This quantity *τ* is the maximum time that the vesicle stays within any 300nm radius sphere. For any vesicle that never moves more than 300nm from its initial simulation location, *τ* = 3600 is assigned. The 300nm threshold is the size of a large insulin vesicle. For each simulation condition, we will compute the distribution of *τ* values over independent and identical simulations.

For a list of all parameters for the discrete model, see Table 2.

**Table 2:** Synopsis of the Model Parameters. For the number and length of filaments, the larger (resp. smaller) number produces a network with 68% (resp. 34%) of spaces <240nm in diameter. These are referenced as dense and sparse networks respectively.

| Parameter Value | Value | Units |
| --- | --- | --- |
| Number of MTs | 15,000 (10,000) | Number |
| Number of vesicles | 200 | Number |
| Cell size | 10 | $\mu m$ |
| MT length | 7 (4) | $\mu m$ |
| Diffusion coefficient | 0.01 | $\mu m^2/s$ |

## RESULTS

In these results we assess how properties of the filament network, the size of the vesicle, and the interaction between the two influence the mobility of vesicles. Toward this end, we first assess what fraction of the space in a fixed network (MT half-life = ∞) is available to vesicles of different sizes (diameter). To do this, a Monte Carlo approach (see Figure 2 caption) is used to determine the fraction of randomly chosen locations in a cell that permit the placement of vesicles of different sizes. This illustrates that for vesicles larger than >200nm in diameter, only a small portion of the volume of the cell is accessible in either the dense or sparse network cases. This is important since these are common insulin vesicle sizes.

**Figure 2.**
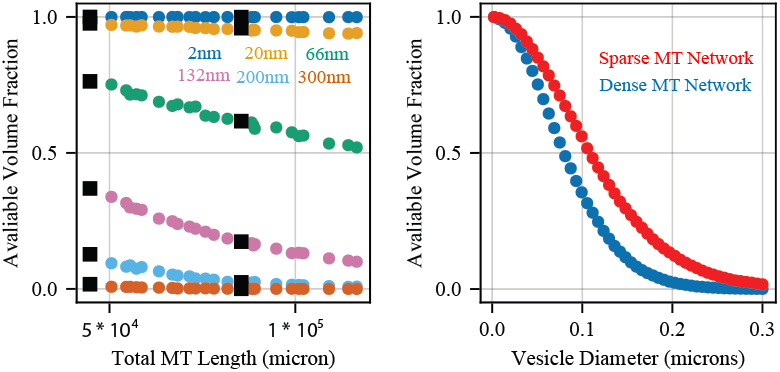
(Color online) Left: We simulate networks with different values of (N,L) as shown in Table 1. For each network, the total length of MT in the cell is computed. For each network 1,000 points are randomly seeded in the cell at least 1 *μ*m from the membrane and the distance to the closest MT is computed. For vesicles of different sizes (given in diameter), the fraction of those distances that permit the placement of a particle of that size is computed. This fraction is plotted as a function of the total MT length in that network. The two sets of black squares indicate the “dense” and “sparse” networks discussed in (Table 1). Right: The same quantity is plotted for the dense and sparse networks as a function of vesicle diameter.

We note this analysis does not account for the potential for vesicles to deform filaments or be deformed by them. Full accounting of this would dramatically increase compute complexity and time. Thus, this available volume fraction should be viewed as a lower limit.

### Analysis of vesicle transit to membrane

Glucose simulated insulin secretion (GSIS) requires the transit of insulin vesicles from the interior of the cell to the near membrane region (defined as 200nm or closer here). After stimulation, *β* undergo a primary secretion phase of ~ 5min followed by a secondary phase lasting up to 60min (34). Here we quantify what fraction of the 200 independent simulations reach the cell membrane within 60 simulated minutes (Table 3). Results show the following. Vesicles <200nm in size are mostly unhindered by the filament network in all cases. The likelihood of a larger vesicle reaching the membrane is however significantly reduced and significantly depends on the dynamics and density of the network. The reduction in density from the dense to the sparse network significantly increases the fraction of vesicles that reach the membrane. Further, a decrease in the MT half-life (faster dynamics) also increases fraction that reach the membrane. These results suggest that at practical densities, normally sized insulin vesicles are significantly constrained by the filament network. Further, dynamics of that network can reduce this constraint and improve insulin availability. Glucose stimulated increase in MT dynamics (observed in (13)) thus provides a mechanism to increase the availability of readily releasable insulin when it is needed.

**Table 3:** Membrane passage fraction. Fraction of particles that reach the boundary in 60 minutes in MT networks of different filament lifetimes. The values are calculated for both a “dense” MT network with *L* = 7, *N* = 15, 000 (left of ‘/’) and a “sparse” MT network with *L* = 4, *N* = 10, 000 (right of ‘/’).

| MT life / Vesicle diameter | 2nm | 20nm | 66nm | 132nm | 200nm | 300nm |
| --- | --- | --- | --- | --- | --- | --- |
| 10 sec | 1.0/1.0 | 1.0/1.0 | 1.0/1.0 | 0.99/1.0 | 0.69/0.95 | 0.12/0.72 |
| 100 sec | 1.0/1.0 | 1.0/1.0 | 1.0/1.0 | 0.91/1.0 | 0.47/0.88 | 0.02/0.60 |
| 1,000 sec | 1.0/1.0 | 1.0/1.0 | 1.0/1.0 | 0.90/1.0 | 0.28/0.80 | 0.00/0.36 |
| 10,000 sec | 1.0/1.0 | 1.0/1.0 | 1.0/1.0 | 0.82/0.96 | 0.15/0.74 | 0.00/0.34 |
| Fixed MT | 1.0/1.0 | 1.0/1.0 | 1.0/1.0 | 0.84/0.96 | 0.20/0.80 | 0.00/0.34 |

### Network density slows vesicle motions while network dynamics induce ‘anomalous’ motions

The previous section suggests the network significantly restrains the motions of larger vesicles. Here we analyze this effect in more detail by analyzing the statistical properties of the vesicle motions. We first quantify the TAMSD as a function of *s* (see Methods) for each independent simulation in each condition. Figure 3 demonstrates that vesicles <200nm in diameter appear to be diffusive (Figure 4 shows similar for the more sparse network). This suggests the network doesn’t substantially restrict the motion of these vesicles, consistent with the prior analysis of membrane passage in Table 3.

**Figure 3.**
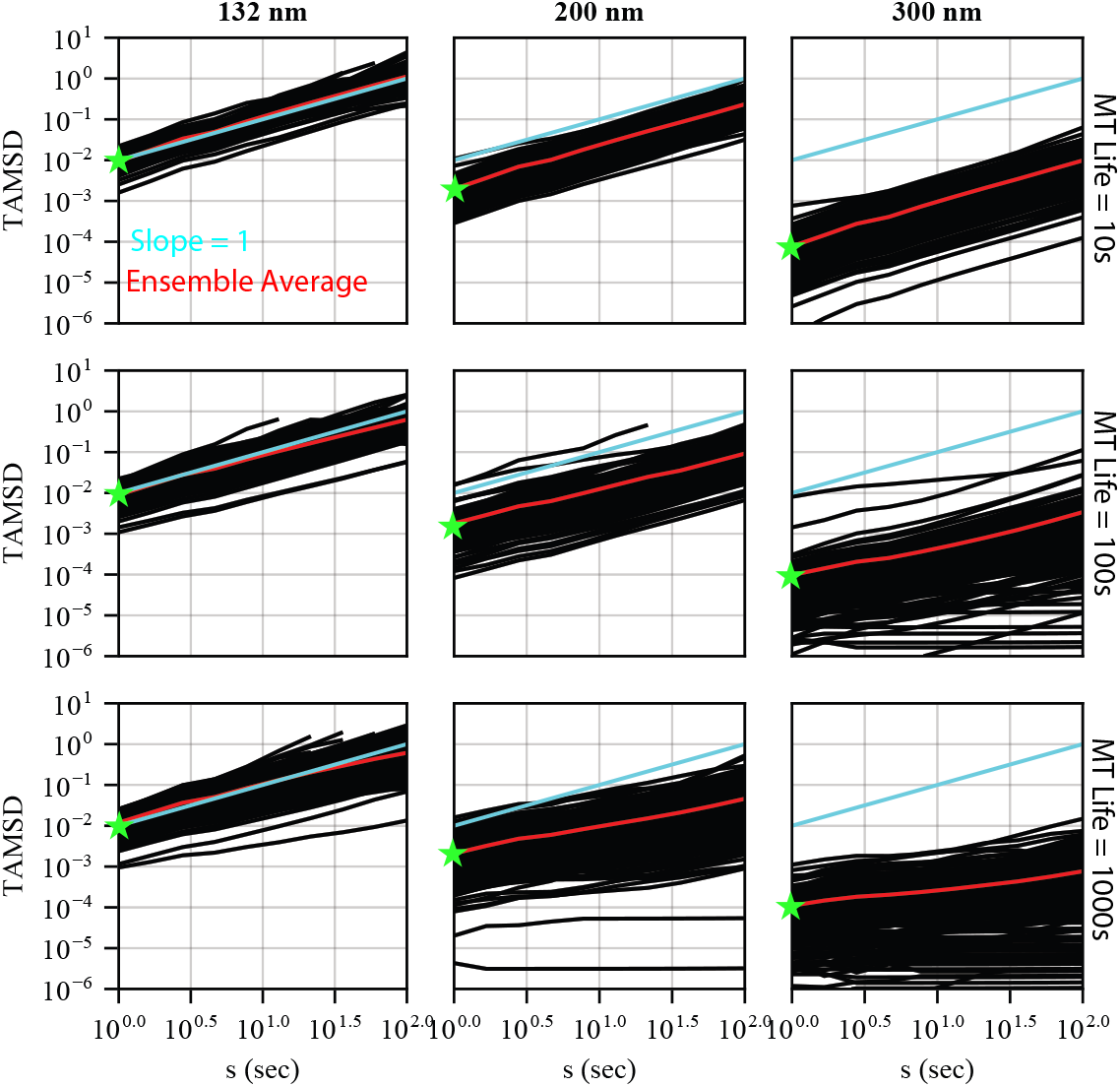
(Color online) TAMSD versus *s* for particles of different size with the L=7, N=15,000 “Dense” network. 200 particles of each size were simulated independently. Red lines show the average TAMSD over the 200 particles. Blue lines are reference lines of slope 1 (standard diffusion) in log-log space for comparison. Slopes of the red lines are given in Table 4. The green stars indicate the *t* = 1 value for the red curves which serve as estimates for the effective diffusion coefficient (recall *MSD* ~ *D*_*s*_^*α*^). For reference, the free space diffusion coefficient used in all cases is *D* = 10^−2^. Smaller particles with diameter 2, 20, 66 nm were also simulated but demonstrated standard diffusion characteristics similar to the top left panel. Results show that larger particles exhibit significant deviations from standard diffusion.

**Figure 4.**
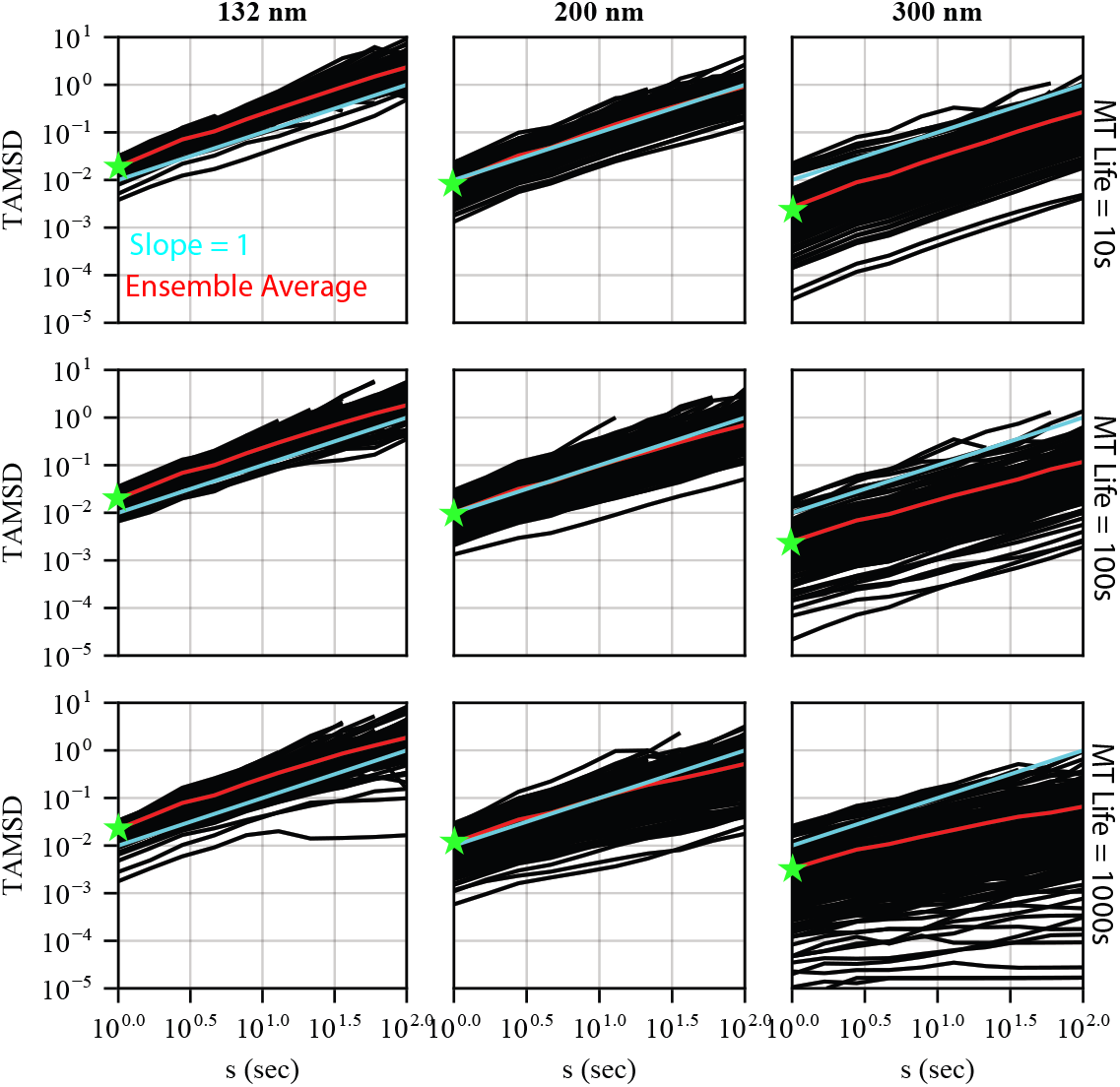
(Color online) The same as Figure 3 but with the sparse MT network with L=4, N=10,000.

For larger vesicles however we see a significant spreading of the trajectories with a significant shallowing of the slope (<1) for some trajectories. This is further illustrated by the deviation of the ensemble averages (red lines) from a slope of one. Table 4 collects the slopes of these ensemble averages in different conditions for comparison. Here, the slope of the ensemble average (over 200 independent simulations) TAMSD is shown for vesicles of different size in networks of different densities and speeds of remodeling. Larger vesicles exhibit deviation from a standard diffusion scaling, this effect is more prominent in the more dense network, and the effect reduces as the network becomes more dynamic (shorter MT half-life).

**Table 4:** TAMSD slope. Slope of the TAMSD versus (*s*) as a function of vesicle diameter and MT half-life. This is calculated for both a “dense” MT network with *L* = 7, *N* = 15, 000 (left of ‘/’) and a “sparse” MT network with *L* = 4, *N* = 10, 000 (right of ‘/’).

| MT life / Vesicle diameter | 2nm | 20nm | 66nm | 132nm | 200nm | 300nm |
| --- | --- | --- | --- | --- | --- | --- |
| 10 sec | 1.0/1.0 | 1.0/1.0 | 1.0/1.0 | 1.0/1.0 | 1.0/1.0 | 1.0/1.0 |
| 100 sec | 1.0/1.0 | 1.0/1.0 | 1.0/1.0 | 0.92/0.98 | 0.83/0.90 | 0.62/0.79 |
| 1,000 sec | 1.0/1.0 | 1.0/1.0 | 1.0/1.0 | 0.85/0.96 | 0.60/0.84 | 0.37/0.64 |
| 10,000 sec | 1.0/1.0 | 1.1/1.0 | 1.0/1.0 | 0.84/0.96 | 0.53/0.84 | 0.27/0.54 |
| Fixed MT | 1.0/1.0 | 1.0/1.0 | 1.0/1.0 | 0.85/0.97 | 0.48/0.80 | 0.22/0.53 |

Figures 3, 4 further show the effect of MT density and dynamics on the effective diffusion constant. In the case of anomalous diffusion *MSD* = *Dt* ^*α*^. Thus the intercepts of the MSD curves at *t* = 1 are estimates of *D* in each condition. In the case of the dense network, *D* reduces by 100x between 132nm and 300nm vesicle sizes (for all MT half-lives). In the case of the sparse network (Figure 4), this reduction is only ~ 5 − 10*x* for all MT half-lives.

Thus we see two separate effects of the filament network on vesicle dynamics. 1) The network generally slows the motions of vesicles, as illustrated by the reduction in *D*. This effect is dependent on the interaction between vesicle size and network density, but not network dynamics. 2) The network introduces an anomalous nature to the vesicle dynamics. This is illustrated by the deviation of the log-log slopes from 1. This effect is stronger for larger vesicles in more dense networks. However this effect is reduced significantly when network dynamics are faster.

The net effect of these two influences is a significant restriction in the mobility of insulin vesicles. To further illustrate this, we quantified the duration of periods during which each vesicle is effectively “immobile”. Here we define immobile as any period during which the vesicle exhibits a displacement of < 300*nm* (see Methods for further detail). For each vesicle path, we find the longest such duration and plot the distribution of these duration over 200 independent simulations (Figure 5). The 66nm vesicle exhibits simple diffusion under all MT conditions (results above) and its distribution in Figure 5 is a point of comparison. Note also that total simulation time is 3600 sec. Thus any distribution mode >3000 sec represents a vesicle that effectively never deviates more than 300nm from its initial location. Beginning at 132 nm we begin to see that a fraction of the 200 simulated vesicles begin to exhibit some period of significant restriction. This is illustrated by the appearance of distribution mass >1000 sec as well as the appearance of a mode >3000 sec. At 132nm, this is most prominent for the dense network with slower dynamics. This appearance of restriction becomes more prominent with larger vesicle diameters. In the slowest network (MT Life = 1000 sec), few of the simulated vesicles leave their initial location.

**Figure 5.**
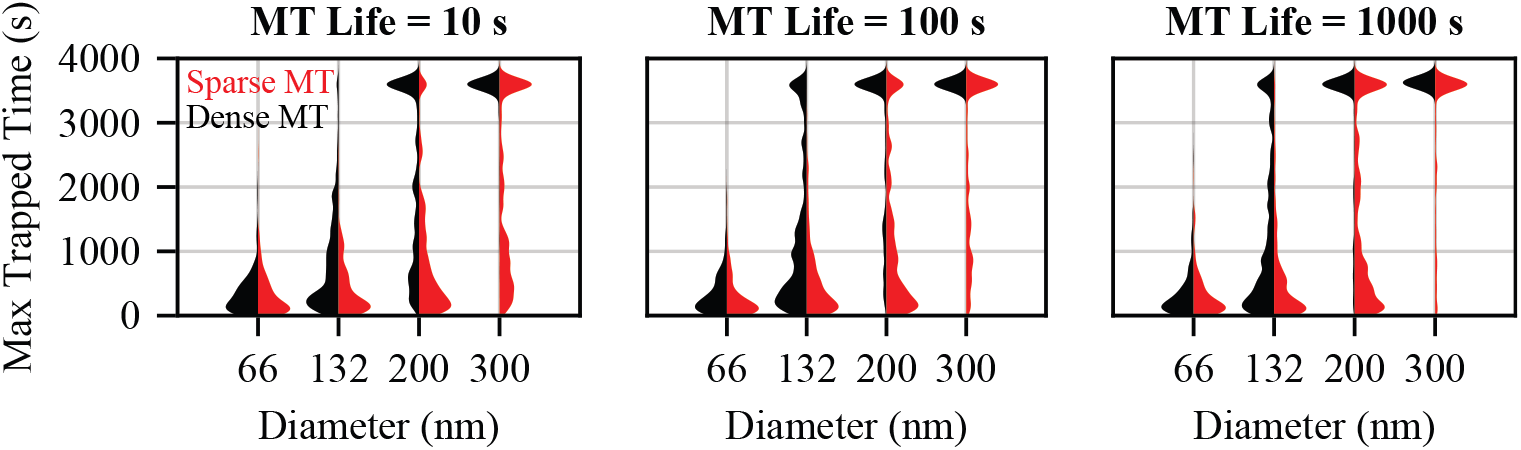
(Color online) Here we assess the extent to which vesicles of different size are constrained by networks of different density and dynamics. For each simulated trajectory, we find the maximum duration during which the vesicle stays within a 300nm radius bubble. We then plot a violin plot over the 200 identical and independent trajectories for each condition to assess how the distribution of this quantity changes with vesicle size and MT dynamics. Note that the maximum simulation time is 3600 sec. Times beyond that are an artifact of the violin plotting. Any mode above 3000 sec is a vesicle that effectively never moves during the entire 3600sec trajectory. Vesicles of diameter 2, 20nm are also simulated but results are similar to the 66nm case. Longer MT lifetimes (10,000 sec) and a fixed MT network are also simulated but show similar results as the 1,000 sec case.

### Dynamic network remodeling causes anomalous mobility via dynamic caging

The previous results suggest that, in some network conditions, larger vesicles rarely displace >300nm from their initial location (Figure 5). There are two possible reasons for this. The first is that the vesicle becomes “caged” by the filaments around it. The second is that the vesicle motions are simply slowed to the point that you would not statistically expect the vesicle to move further than this due to slowed motions.

We thus next look for signatures of “caging” or CTRW behavior in the simulated vesicle tracks by quantifying TAMSD as a function of *t* (Figure 6). For vesicles < 132*nm*, these curves are all flat indicating ergodic behavior. For larger vesicles however, some of these curves slope downward as a function of *t*. This is indicative of an “aging” phenomenon where longer periods of immobility are found when longer runs of track data (*t*) are included in the analysis. This is a breaking of ergodicity and a hallmark of CTRW behavior. Interestingly this aging phenomenon is most prominent when the MT dynamics are sufficiently fast (lower MT half-life).

**Figure 6.**
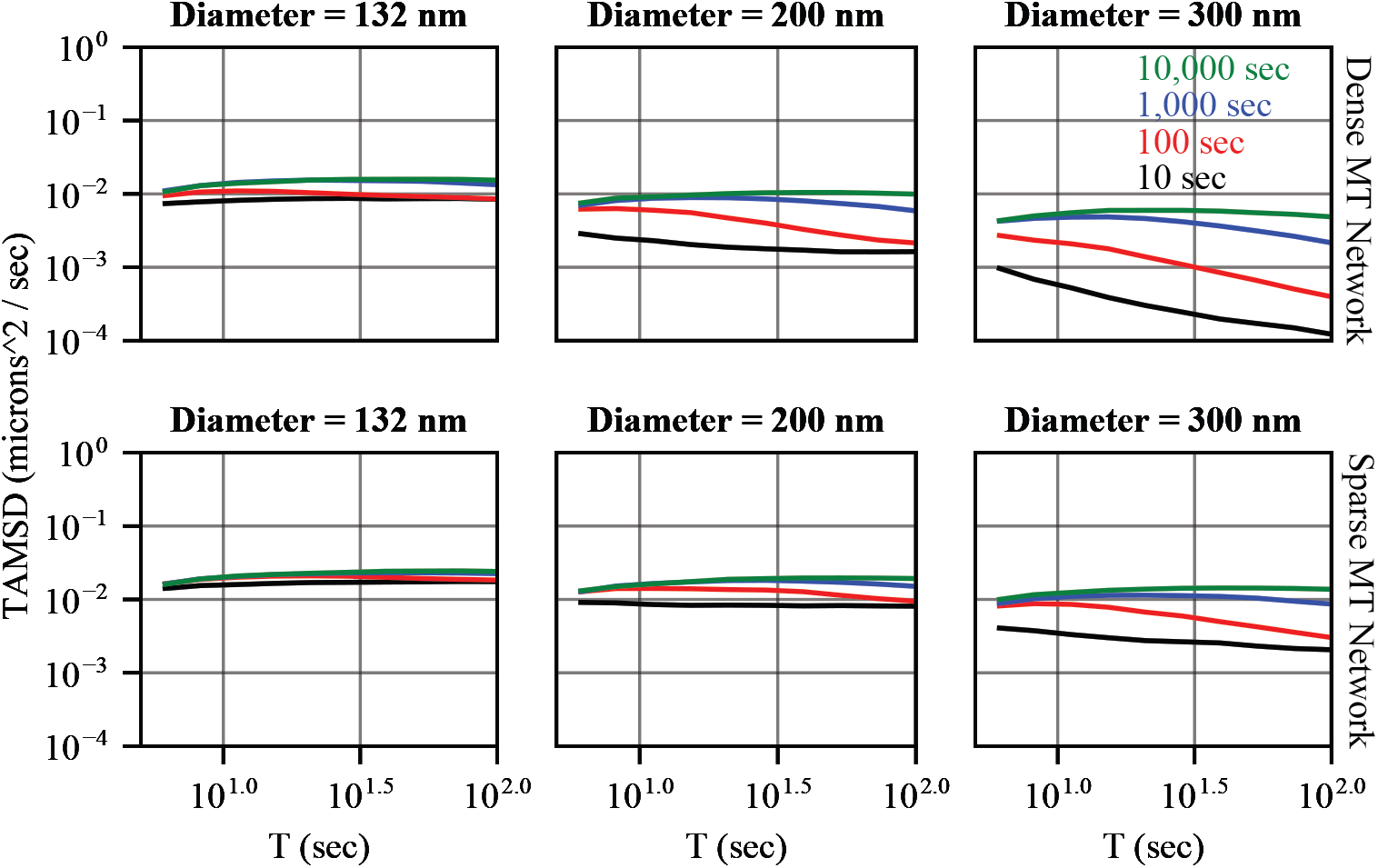
(Color online) Simulation of the effect of the speed of MT dynamics on ‘aging’. TAMSD is plotted for different observation window lengths (T = 5 - 100), with S=5 sec. The legend colors indicate the mean lifetime of each MT in seconds. The top row shows results for the “dense” MT network (L=7, N=15,000) and the lower row for the “sparse” network (L=4, N=10,000). A downward-sloping line indicates “aeging”, the phenomenon seen by Taberi et al. This effect becomes apparent for sufficiently fast MT dynamics and sufficiently large particles and is more pronounced for the Dense network than the sparse.

We posit the following explanation for this observation. Sufficient density of MTs is clearly needed to create the conditions that immobilize these vesicles. However, in a static network any individual vesicle is either caged or not. The dynamics of the network are responsible for remodeling the environment, freeing some vesicles from their cages and enclosing others. The longer the duration of data that is analyzed, the higher the chance that one of those cages will persists for a longer period, leading to a decrease in TAMSD as a function of *t*. This suggests that the statistical observation of anomalous motions of insulin are a consequence of the dynamics of these MTs. Further, this anomalousness becomes more prominent as the density of the network increases relative to the size of vesicles.

### Study limitations

There are two main areas of limitations in the model we construct. Fist, the physical structure of the cell is more complex and packed environment than we account for here. We do not account for the presence of organelles and other structures that occupy space in the cell. MTs are fixed in place (though they can be de polymerized and new MTs nucleated) and thus we do not model the steric interactions between filaments. Lastly, we do not model the interactions between different vesicles or the deform-ability of vesicles in the cell. Our main focus is on how the MT network influences mobility and so that is the main focus of the modeling. These limitations can likely be overcome, though at the expense of a drastic increase in computational complexity and time.

The second main area of limitations is our treatment of the MT dynamics. There are numerous potential sources of dynamics including dynamic instability (35), depolymerization of existing and nucleation of new filaments, and motor driven transport of MTs along other MTs (36). MTs are generally stable in *β* cells and thus we do not incorporate dynamic instability (13). However motor-driven MT sliding is common (36) and likely a substantial source of network dynamics. With *N* = 10, 000 − 15, 000 MTs, each interacting with potentially multiple other MTs in a disordered network, accounting for this quickly becomes intractable. We thus use the destruction and construction of MTs as a general proxy for dynamics. MT sliding could reasonably have similar effects on insulin mobility that we find in this study, though this would need to be studied further.

## CONCLUSION AND DISCUSSION

The goal of this study was to determine how the dense meshwork of MT filaments influences the mobility and availability (for GSIS) of insulin vesicles. Toward this end, we constructed a computational model of the MT network in insulin-producing *β*-cells to investigate insulin vesicle transport dynamics. To mimic the structure of the MT network in actual *β* cells, we calibrated the number (N) and length (L) of the filaments to match the statistical density observed in measured cells. We further incorporated MT dynamics through random depolymerization and nucleation events, with varying half-lives to reflect different levels of dynamics of the network. Using this model we explored how vesicles interact with the MT network and assessed the impact of network properties on vesicle mobility and availability for GSIS.

Our results highlight the significant effects the MT network has in regulating insulin granule transport and availability for secretion in pancreatic *β*-cells. Both network density and dynamics critically influence the mobility of insulin within the cell and its availability (ability to reach the membrane). When the MT network density is calibrated to that observed in imaged *β* cells, vesicles smaller than 200nm in diameter appear to be largely unhindered by the MT network. Their movement closely resembles standard diffusion and the presence of the MTs does not significantly reduce their ability to reach the cell membrane. This is consistent with prior studies suggesting that smaller vesicles can move relatively freely in cellular environments (37–41).

However, insulin vesicles commonly range from 100-300nm in diameter and our results indicate mobility of larger vesicles is significantly impaired by the MT network. The network slows the speed of larger vesicles (>200nm) motion (as measured by a generalized diffusion coefficient), leads to their confinement in MT “cages”, and induces anomalous diffusion behavior. These effects depend on both the density of the network and on the speed of its dynamical remodeling. As the network becomes more dense, the restrictions it imposes on vesicle motions becomes more significant. While this is conceptually obvious, our results demonstrate that this restriction is substantial for normal-sized vesicles (200-300nm) in MT networks at densities observed in *β* cells.

The dynamics of the MT network also have an influence on vesicle mobility. At a fixed density, vesicles are more mobile in more dynamic networks. This increased mobility has the effect of increasing the likelihood of vesicles reaching the cell membrane. Thus, increased speed of network dynamics can increase the size of the available pool of readily releasable vesicles, at least in the sense of increasing the fraction that can physically reach the membrane (we do not consider modification of the vesicles for docking and exocytosis). Detailed analysis of the TAMSD of simulated vesicles reveals that this increased mobility is not due to increased speed of motions (as measured by a generalized diffusion coefficient). Speed of motion depends on network density but not dynamics. Instead, dense networks cage large vesicles and dynamics of the network lead to cyclical caging and un-caging. The dynamic un-caging increases mobility by ensuring vesicles are not completely immobilized.

The network dynamics also introduce anomalous behavior in the motions of vesicles. Specifically, it introduces an “aging” behavior for larger vesicles which is characteristic of anomalous sub-diffusion. Interestingly, this effect only occurs when the network density is sufficiently high (relative to the size of the vesicle) but is not the result of the simple presence of a dense network. Instead, the dynamics of the network induce this anomalous behavior. We posit the following explanation for this. Aging is a consequence of the vesicles becoming periodically caged, with longer durations of immobility becoming more likely with longer observation windows. In a static network, a vesicle is either caged and immobile or un-caged and mobile. It is the dynamics of the network that induces the cycling between these two states, and thus the anomalous motion.

Taken together, these results suggest that MTs, at physiological densities, substantially restrict the mobility and availability of insulin for GSIS. Further, this effect is weakened when the MT network displays increased speed of dynamic remodeling.

This is consistent with a number of prior experimental observations. 1) Depolymerization of MTs by nocodazole substantially increases GSIS while their stabilization by taxol substantially decreases GSIS (13). 2) Increased MT density correlates with decreased GSIS and diabetic mouse models exhibit higher density of MTs (13). 3) Glucose stimulation increases the speed of MT depolymerization and nucleation (13), potentially increasing availability of insulin when it is needed. 4) Glucose stimulation additionally increases the rate of motor-driven MT sliding (36), an alternative form of dynamics that might serve to further free vesicle mobility. 5) The aging phenomenon that is a consequence of MT dynamics was previously observed for insulin vesicle transport in *β* cells.

These results add to a growing body of research suggesting there is potential therapeutic merit in targeting the MT cytoskeleton to modulate insulin availability for GSIS. While gross manipulation of the cytoskeleton may come with undesirable side effects, these results indicate the potential for targeted manipulations. MT (de)stabalizers could be applied to up / down regulate GSIS by modulating density. Alternatively, dynamics could be altered without substantially affecting density. Glucose stimulation already speeds up MT dynamics by speeding de polymerization, nucleation, and sliding. These processes could be targeted pharmacologically to elicit desired changes in GSIS. This is of course speculation at this stage. However these and prior results do demonstrate that MTs substantially influence insulin mobility and availability, and thus further research is needed to further detail this influence and determine its suitability as a therapeutic target.

## AUTHOR CONTRIBUTIONS

TT and WRH jointly participated in the conceptualization and design of this work, carried out the work, and wrote this article.

## ACKNOWLEDGMENTS

This work was supported by NIH grant 2R01 DK106228.

## SUPPLEMENTARY MATERIAL

An online supplement to this article can be found by visiting BJ Online at http://www.biophysj.org.

